# Fluorochrome-dependent specific changes in spectral profiles using different compensation beads or human cells in full spectrum cytometry

**DOI:** 10.1101/2023.06.14.544540

**Authors:** Yaroslava Shevchenko, Isabella Lurje, Frank Tacke, Linda Hammerich

**Author notes:** corresponding author: Dr. rer. nat. Linda Hammerich, Department of Hepatology and Gastroenterology, Charité Universitaetsmedizin Berlin, Augustenburger Platz 1, 13353 Berlin, Germany. **Ethics statement:** The study was conducted according to the guidelines of the Declaration of Helsinki, and approved by the local ethics committee at Charité Universitätsmedizin Berlin, Germany (ethical approval number EA2/065/21). Informed consent was obtained from all subjects involved in the study and is kept on file.

## Abstract

Full spectrum flow cytometry is a powerful tool for immune monitoring on a single-cell level and with currently available machines, panels of 40 or more markers per sample are possible. However, with an increased panel size, spectral unmixing issues arise, and appropriate single stain reference controls are required for accurate experimental results and to avoid unmixing errors. In contrast to conventional flow cytometry, full spectrum flow cytometry takes into account even minor differences in spectral signatures and requires the full spectrum of each fluorochrome to be identical in the reference control and the fully stained sample to ensure accurate and reliable results. In general, using the cells of interest is considered optimal, but certain markers may not be expressed at sufficient levels to generate a reliable positive control. In this case, compensation beads show some significant advantages as they bind a consistent amount of antibody independent of its specificity. In this study, we evaluated two types of manufactured compensation beads for use as reference controls for full spectrum cytometry and compared them to human and murine primary leukocytes. While most fluorochromes show the same spectral profile on beads and cells, we demonstrate that specific fluorochromes show a significantly different spectral profile depending on which type of compensation beads is used, and some fluorochromes should be used on cells exclusively. Finally, we provide a list of appropriate reference controls for 30 of the most commonly used and commercially available fluorochromes.

## Statement of purpose

Full spectrum flow cytometry is a powerful tool for immune monitoring on a single-cell level with extensive use in cancer immunotherapy research as well as in investigation of various immunological diseases[1, 2]. With the availability of 5-laser instruments and novel fluorophore families, panels of 40 or more markers became possible [3, 4]. However, with an increase of panel size, spectral unmixing issues arise, and panel optimization is becoming increasingly important to maximize the potential data outcome from each specimen[5]. Appropriate single stain reference controls are required for accurate experimental results and to avoid unmixing errors[6]. Most commonly, reference controls are compensation beads or cell samples stained with a single fluorescently-labeled antibody used in the experiment, which provides the spectral signature of each dye for the unmixing algorithm and establishes a baseline for comparison to the studied sample[5, 7].

In contrast to conventional flow cytometry, which solely focuses on the signal within the specific wavelength bandwidth of interest, full spectrum flow cytometry takes into account even minor differences in spectral signatures that can impact the reliability of reference controls. The full spectrum of each fluorochrome must be identical in the reference control and the fully stained sample to insure accurate and reliable results[5]. Otherwise, inappropriate reference controls may lead to a difference in signal intensity and spread and ultimately, unmixing errors[8] . As a result, interpretation of the data might be incorrect due to inappropriate gating, variability in data obtained from different samples and reduced sensitivity in detecting low fluorescence levels.

When establishing reference populations both beads and cells have proven to be valuable options for single stain controls. In general, using the same type of cells that will also be analyzed in the experiment is considered optimal, since this takes into account the autofluorescence profile of the cells of interest[9]. However, certain markers may not be expressed at sufficient levels or on enough cells to generate a reliable positive control (for example, lineage markers for dendritic cells, which represent less than 1% of leukocytes in the peripheral blood[10]). In this case, compensation beads show some significant advantages as they bind a consistent amount of antibody independent of its specificity[11]. In addition, they are commercially available and relatively inexpensive ready-to-use products, while the availability of biological specimens, especially when working with human samples, might be limited.

In this study, we evaluated two types of manufactured compensation beads for use as reference controls for full spectrum cytometry and compared them to primary leukocytes from both human and mouse. We provide a list of appropriate reference controls for 30 of the most commonly used and commercially available fluorochromes. While most fluorochromes show the same spectral profile on beads and cells, we demonstrate that specific fluorochromes show a significantly different spectral profile depending on which type of compensation beads is used, including some that should be used on cells exclusively.

## Materials and Methods

### Single stains on compensation beads

One drop of ThermoFisher Ultracomp eBeads Plus™ (Thermo Fisher Scientific, Waltham, USA) or one drop each of positive and negative Biolegend compensation beads (Biolegend, San Diego, USA) were diluted in 1ml of staining buffer (HBSS +2mM EDTA). After thorough vortexing 100μl of the diluted compensation beads were used for each reference control. Fluorochrome-conjugated antibodies were added at the indicated concentrations (Supplementary Table 1) and incubated for 15 min at room temperature. A washing step was performed with 2ml of staining buffer (HBSS +2mM EDTA) and samples were resuspended in 200μl of staining buffer for acquisition.

### Single stains on human peripheral blood leukocytes

Anti-coagulated (EDTA) whole blood was obtained from volunteers (ethical approval number EA2/065/21) and processed immediately. 200μl of whole blood were incubated with indicated concentrations (Supplementary Table 1) of fluorochrome-conjugated antibodies for 20 min at room temperature. Lysis of red blood cells was performed by adding 2ml of 1x BD Pharm Lyse™ Lysing buffer and incubating the samples for 10 min at room temperature. Cells were washed in staining buffer (HBSS +2mM EDTA) and incubated with lysis buffer a second time for 5 min at room temperature. After washing, cells were fixed in 2%formalin (in 1x PBS) for 10 min. After the last washing step, cells were resuspended in 200μl staining buffer to proceed with acquisition.

### Single stains on mouse splenocytes

Mouse splenocytes were harvested by forcing a spleen from a wildtype mouse through a 70μm nylon mesh to achieve a single-cell suspension. Pelleted cells were resuspended in 5ml of 1x BD Pharm Lyse™ Lysing buffer and incubated for 5 min at room temperature to remove red blood cells. Cells were washed with HBSS and resuspended in an appropriate volume of staining buffer. After filtering through a 70μm mesh again, 10% of the single cell suspension were used for each reference control. Fluorochrome-conjugated antibodies were added directly to the cells at the indicated concentration (Supplementary Table 1) and incubated for 20min at room temperature.

### Data acquisition and analysis

Sample analysis was performed on a standard issue Cytek® Aurora Cytometer equipped with 3 lasers and 38 detectors (Supplementary Table 2) and running Spectroflo® v3.0.3 (Cytek Biosciences, Inc.). QC was performed daily, and samples were acquired at medium speed using Cytek Assay Settings. The stopping gate was set at 2,500 events for beads or 100,000 events for cells. FSC and SSC were adjusted individually depending on the nature of the single stain control: compensation beads, human or mouse immune cells.

### Normalized spectra and similarity

Normalized spectra for each reference control were extracted from Spectroflo® v3.0.3 as follows: Interval gates for positive and negative peaks were defined for each single stain in the respective peak channel (Supplementary Table 3) and statistics tables indicating the median signal intensity of all fluorescent channels were created for both the positive (MFI_pos_) and the negative (MFI_neg_) gate. Statistics data was exported to Excel, and the normalized signal intensity for each fluorescent dye in each channel was calculated as MFI_pos_-MFI_neg_/(MFI_pos_-MFI_neg_)max. Using normalized medium fluorescence intensity data, spectrum overlays were created in GraphPad Prism 9 and visualized as a histogram overlay graph for comparison. The similarity between indicated fluorochromes was determined by the unmixing tool of SpectroFlo® and displayed in heat maps, values inside the cells are given in %.

## Results

The availability and diversity of manufactured compensation beads have significantly increased in the past years as they are increasingly used as reliable reference controls in flow cytometry. However, not all types of compensation beads bind all antibody species (Supplementary Table 4), which reduces their convenience of use. We compared the species reactivity of commercially available compensation beads (Supplemetary Table 4) and decided to test Biolegend Compensation beads as well as ThermoFisher Ultracomp eBeads™ Plus due to their ability to bind the most indicated species. We tested 30 of the most commonly used and commercially available fluorochromes in full spectrum cytometry on both types of beads, as well as human peripheral blood leukocytes and mouse splenocytes. The list of all antibodies used in this study can be found in Supplementary Table 1. In general, we used the same anti-human antibody on both types of beads as well as human peripheral blood leukocytes (Supplementary table 3), while a different antibody was used on murine splenocytes due to specificity.

25 out of the 30 tested fluorochromes showed identical emission spectra on cells and either type of beads (Supplementary Figure 1). Consistently, Spectrflo^®^ determined a similarity of 100% for all reference control types for each of these fluorochromes (Supplementary Figure 2). This enables the use of compensation beads as reliable reference controls for each tested color.

However, five fluorochromes (BV480, BV785, PerCP-Cy5.5, PerCP-eFluor710, AlexaFluor700) showed different emission spectra on cells and beads (Figure 1). BV480, BV785 and PerCP-eFluor710 showed identical emission spectra on mouse and human primary cells as well as Biolegend Compensation Beads, but different spectra when used on ThermoFisher UltraComp eBeads Plus (Figure 1A, B, D). This is also reflected in the similarity matrices generated by SpectroFlo^®^. PerCP-Cy5.5 showed different emission spectra between all types of primary cells as well as compensation beads (Figure 1C). While primary mouse cells seem to show a spectrum identical to Biolegend Compensation Beads and almost identical to human cells (similarity 99%), UltraComp eBeads™ Plus only reach a similarity of 92 or 94% with mouse and human primary cells, respectively. Notably, the spectra for AlexaFluor 700 seemed to differ between murine cells and Biolegend Compensation Beads vs. human cells and UltraComp eBeads Plus, but only the difference between human and murine cells is picked up by SpectroFlo^®^ (Figure 1E). These results indicate a remarkable advantage of Biolegend compensation beads as a reliable reference control for spectral flow cytometry for all tested fluorochromes except PerCP-Cy5.5.

**Figure 1.**
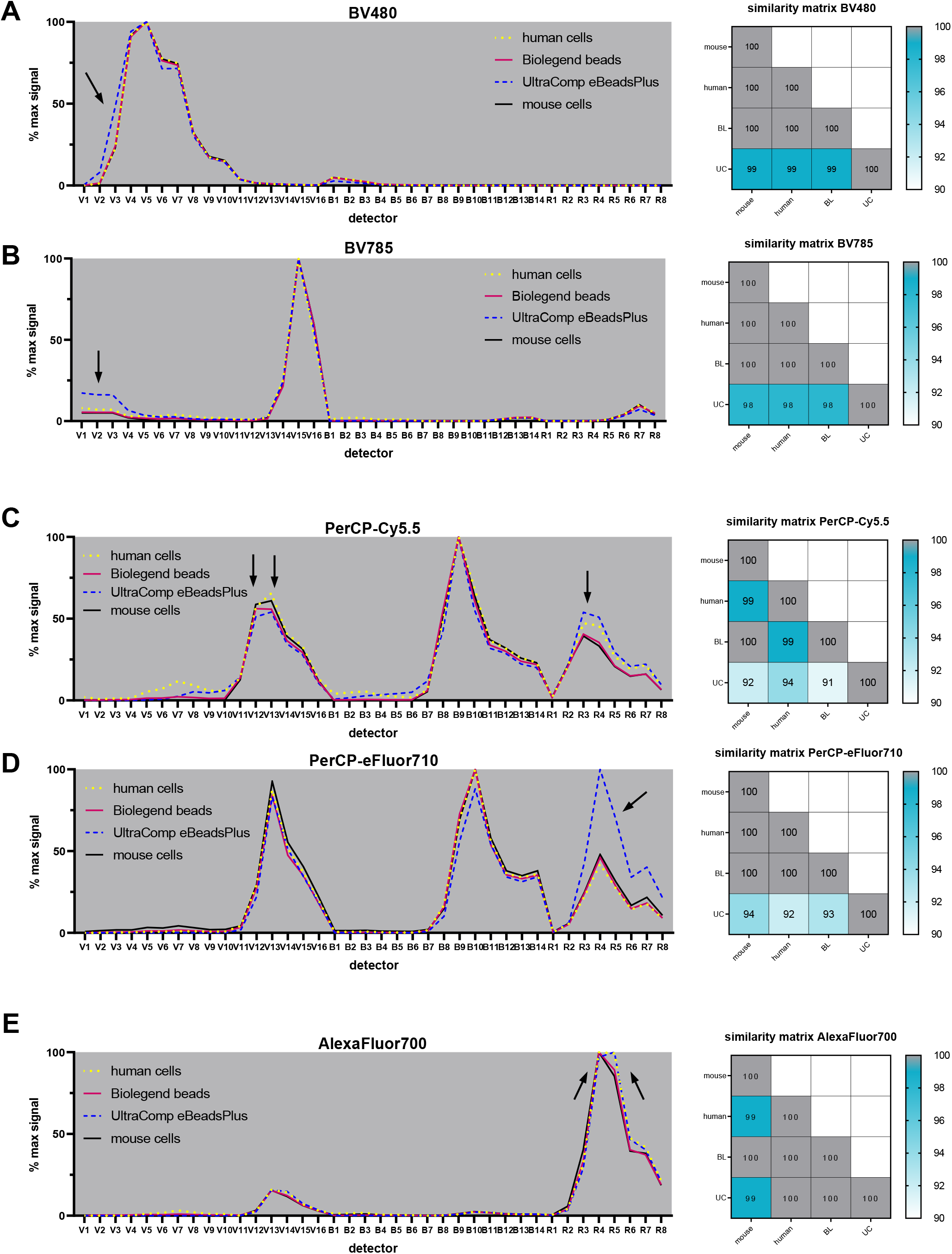
Normalized emission spectra and similarity matrizes of fluorochromes showing different spectra on cells and beads. Indicated fluorochromes were stained on cells (human peripheral blood leukocytes or murine splenocytes), Biolegend® Compensation Beads or UltraComp eBeads™ Plus Compensation Beads. Emission spectra were extracted from Spectroflo software, normalized to the maximum signal and overlaid in GraphPad Prism. Similarity was determined in Spectroflo software. **A** BV480 **B** BV785 **C** PerCP-Cy5.5 **D** PerCP-eFluor710 **E** AlexaFluor 700. BL= Biolegend compensation beads, UC= UltraComp eBeads Plus

Since several of the fluorochromes described in Figure 1 are tandem dyes (BV785, PerCP-Cy5.5, PerCP-eFuor710), we next investigated the potential impact of different antibodies (with different specificity) on the spectral profiles. To test this, we stained both types of compensation beads with several antibodies coupled to the same fluorochromes that showed differential emission spectra before. Supplementary Figure 3 provides emission spectra and similarity matrices for each indicated fluorochrome, the spectrum observed on human cells was kept as a reference. This analysis confirmed a reproducible pattern of similarity as observed in Figure 1, independent of the antibody used.

## Conclusions

In this study, we demonstrate the usefulness and reliability of compensation beads as reference controls for full spectrum cytometry, offering the potential to replace primary cells. However, we also show that the spectral signatures of some fluorochromes differ significantly from cells. Therefore, it is imperative to ensure the emission spectra of the cells of interest are identical to the type of reference controls used for every fluorochrome. Furthermore, our data indicate that the spectral signatures of fluorescent probes are independent of antibody specificity, thereby negating the need for individual verification of each new antibody panel if the fluorochromes and cells of interest are the same. Finally, we provide a readily accessible list (Supplementary table 5) of optimal reference controls for 30 of the most commonly used and commercially available fluorochromes in full spectrum cytometry.

## Acknowledgments

We thank Désirée Kunkel and the BIH Cytometry Core Facility for technical support and helpful discussions.

## Notes

**Conflict of interest disclosure:** The authors declare no financial conflicts of interest.

**Funding:** This work was supported by the German Research Foundation (DFG; CRC1382, SFB/TRR 296 and SPP2306), the Else-Kröner-Fresenius-Stiftung (Grant-ID 2021_EKEA.145) and Charité 3R| Replace - Reduce – Refine.

### Competing Interest Statement

The authors have declared no competing interest.

## References

1. Bonilla, D.L., G. Reinin, and E. Chua, Full Spectrum Flow Cytometry as a Powerful Technology for Cancer Immunotherapy Research. Front Mol Biosci, 2020. 7: p. 612801.

2. Hammerich, L., et al., Resolving 31 colors on a standard 3-laser full spectrum flow cytometer for immune monitoring of human blood samples. Cytometry B Clin Cytom, 2023.

3. Park, L.M., J. Lannigan, and M.C. Jaimes, OMIP-069: Forty-Color Full Spectrum Flow Cytometry Panel for Deep Immunophenotyping of Major Cell Subsets in Human Peripheral Blood. Cytometry A, 2020. 97(10): p. 1044–1051.

4. Sahir, F., et al., Development of a 43 color panel for the characterization of conventional and unconventional T-cell subsets, B cells, NK cells, monocytes, dendritic cells, and innate lymphoid cells using spectral flow cytometry. Cytometry A, 2020.

5. Ferrer-Font, L., et al., Panel Optimization for High-Dimensional Immunophenotyping Assays Using Full-Spectrum Flow Cytometry. Curr Protoc, 2021. 1(9): p. e222.

6. Ferrer-Font, L., et al., Panel Design and Optimization for High-Dimensional Immunophenotyping Assays Using Spectral Flow Cytometry. Curr Protoc Cytom, 2020. 92(1): p. e70.

7. Maecker, H.T. and J. Trotter, Flow cytometry controls, instrument setup, and the determination of positivity. Cytometry A, 2006. 69(9): p. 1037–42.

8. Farrand, K., et al., Using Full-Spectrum Flow Cytometry to Phenotype Memory T and NKT Cell Subsets with Optimized Tissue-Specific Preparation Protocols. Curr Protoc, 2022. 2(7): p. e482.

9. Hulspas, R., et al., Considerations for the control of background fluorescence in clinical flow cytometry. Cytometry B Clin Cytom, 2009. 76(6): p. 355–64.

10. Haller Hasskamp, J., J.L. Zapas, and E.G. Elias, Dendritic cell counts in the peripheral blood of healthy adults. Am J Hematol, 2005. 78(4): p. 314–5.

11. Roederer, M., Compensation in flow cytometry. Curr Protoc Cytom, 2002. Chapter 1: p. Unit 1 14.

