## Supplementary Material for "Fluorochrome-dependent specific changes in spectral profiles using different compensation beads or human cells in full spectrum cytometry"

| fluorochrome | target species | specificity | clone | company | catalog no. | concentration |
|---|---|---|---|---|---|---|
| BV421 | human | CD19 | HIB19 | BioLegend | 302234 | 1:100 |
| | mouse | CD4 | RM4-5 | BioLegend | 100544 | 1:400 |
| SB436 | human | CD31 | WM-59 | Invitrogen | 62-0319-42 | 1:100 |
| | mouse | CD31 | 390 | Invitrogen | 62-0311-82 | 1:400 |
| eFluor450 | human | CD3 | Okt 03 | BD Bioscience | 48-0037-42 | 1:100 |
| | human/mouse | CD11b | M1/70 | Invitrogen | 48-0112-82 | 1:400 |
| BV480 | human | CD4 | RPA-T4 | BD Biosciences | 746541 | 1:100 |
| | mouse | CD4 | RM4-5 | BD Biosciences | 565634 | 1:400 |
| | mouse | Tim4 | RMT4-54 | BD Bioscience | 746499 | 1:400 |
| | mouse | MerTK | 108928 | BD Bioscience | 747896 | 1:400 |
| V500 | human | CD16 | 3G8 | BD Biosciences | 561394 | 1:100 |
| | mouse | CD45 | 30-F11 | BD Biosciences | 557659 | 1:400 |
| SV538 | human | CD3 | UCHT1 | BioLegend | 300484 | 1:100 |
| | mouse | B220 | RA3-6B2 | BioLegend | 103283 | 1:400 |
| BV570 | human/mouse | CD11b | M1/70 | BioLegend | 101233 | 1:100 |
| BV605 | human | CD38 | HIT2 | BioLegend | 303532 | 1:100 |
| | mouse | CX3CR1 | SA011F11 | BioLegend | 149027 | 1:400 |
| BV650 | human | CD15 | W6D3 | BioLegend | 323033 | 1:100 |
| | mouse | CX3CR1 | SA011F11 | BioLegend | 149033 | 1:400 |
| BV711 | human | CD3 | OKT3 | BioLegend | 317328 | 1:100 |
| | mouse | CD8 | 53-6.7 | Biolegend | 100748 | 1:400 |
| BV750 | human | CD56 | 5.1H11 | BioLegend | 362556 | 1:100 |
| | mouse | TCRb | H57-597 | BD Biosciences | 747006 | 1:400 |
| BV785 | human | TLR2 | 11G7 | BD Biosciences | 742771 | 1:100 |
| | mouse | CD8 | 53-6.7 | Biolegend | 100749 | 1:400 |
| | human | CD141 | M80 | Biolegend | 344115 | 1:100 |
| | mouse | CD206 | C068C2 | Biolegend | 141729 | 1:400 |
| | mouse | CD25 | PC61 | BioLegend | 102051 | 1:400 |
| BB515 | human | CD8 | RPA-T8 | BD Biosciences | 564526 | 1:400 |
| | mouse | CD8 | 53-6.7 | BD Biosciences | 564459 | 1:400 |
| FITC | human/mouse | CD11b | M1/70 | BioLegend | 101206 | 1:100 |
| | mouse | CD4 | RM4-5 | BD Pharmigen | 553047 | 1:400 |
| Alexa532 | human | CD45 | HI30 | Invitrogen | 58-0459-42 | 1:100 |
| | mouse | CD45 | 30-F11 | Invitrogen | 58-0451-82 | 1:400 |
| PE | human | CX3CR1 | 2A9-1 | BD Biosciences | 565796 | 1:100 |
| | mouse | CD4 | RM4-5 | Biolegend | 100512 | 1:400 |
| PE/Dazzle594 | human | CCR2 | K036C2 | BioLegend | 357222 | 1:100 |
| | human/mouse | CD11b | M1/70 | BioLegend | 101255 | 1:400 |
| PE/Fire640 | human | CD45RO | UCHL1 | BioLegend | 304263 | 1:100 |
| | mouse | CD11c | QA18A72 | Biolegend | 161104 | 1:400 |
| PE-Cy5 | human | CD19 | HIB19 | BioLegend | 302210 | 1:100 |
| | mouse | CD19 | eBio 1D3 | Invitrogen | 15-0193-83 | 1:400 |
| BB700 | human | CD127 | HLA-7R-M21 | BD Biosciences | 566398 | 1:100 |
| | mouse | CD11c | N418 | BD Biosciences | 745852 | 1:400 |
| PerCP-Cy5.5 | human | CD14 | M5E2 | BioLegend | 301824 | 1:100 |
| | mouse | CD4 | RM4-5 | Invitrogen | 45-0042-82 | 1:400 |
| | mouse | CD103 | 2EF | Biolegend | 121416 | 1:400 |
| | mouse | CD11c | N418 | Invitrogen | 45-0114-82 | 1:400 |
| PerCP-eFl710 | human | CD45RA | GRT22 | Invitrogen | 46-0468-42 | 1:100 |
| | human | CD11c | 3,9 | Invitrogen | 46-0116-42 | 1:100 |
| | human | LAG3 | 3DS223H | Invitrogen | 46-2239-42 | 1:100 |
| | mouse | CD137 | 17B5 | ThermoFisher | 46-1371-82 | 1:400 |
| | mouse | CD369 | bg1fpj | Invitrogen | 46-5859-82 | 1:400 |
| PE-Cy7 | human | CD161 | HP-3G10 | Invitrogen | 25-1619-42 | 1:100 |
| | mouse | CD8 | 53-6.7 | Invitrogen | 25-0081-82 | 1:400 |
| PE/Fire810 | human | CXCR3 | G025H7 | BioLegend | 353759 | 1:100 |
| | mouse | CD3 | 17A2 | Biolegend | 100277 | 1:400 |
| APC | human | CD14 | M5E2 | BioLegend | 301807 | 1:100 |
| | mouse | CD4 | RM4-5 | Invitrogen | 17-0042-82 | 1:400 |
| Al647 | human | CD66b | G10F5 | Biolegend | 305109 | 1:100 |
| | mouse | CD8 | 53-6.7 | Biolegend | 100724 | 1:400 |
| SparkNIR 685 | human | CD45 | 2D1 | BioLegend | 368551 | 1:100 |
| | mouse | CD8 | 53-6.7 | BioLegend | 100781 | 1:400 |
| Al700 | human | CD19 | HIB19 | BioLegend | 302225 | 1:100 |
| | human | CD127 | A7R34 | Invitrogen | 56-1271-82 | 1:100 |
| | mouse | CD8 | 53-6.7 | BD Biosciences | 557959 | 1:400 |
| | mouse | Ly6G | 1A8 | Biolegend | 127622 | 1:400 |
| APC-Cy7 | human | CD33 | WM53 | BioLegend | 303442 | 1:100 |
| | mouse | CD45 | 30-F11 | BD Biosciences | 557659 | 1:400 |
| APC/Fire810 | human | CD25 | M-A251 | BioLegend | 356150 | 1:100 |
| | mouse | Gr1 | RB6-8C5 | Biolegend | 108469 | 1:400 |

**Suppl. Table 1. Antibodies used in this study**

| Laser | Detector | center Wavelength (nm) | bandwidth (nm) |
|---|---|---|---|
| Violet (405nm) | V1 | 428 | 15 |
| | V2 | 443 | 15 |
| | V3 | 458 | 15 |
| | V4 | 473 | 15 |
| | V5 | 508 | 20 |
| | V6 | 525 | 17 |
| | V7 | 542 | 17 |
| | V8 | 581 | 19 |
| | V9 | 598 | 20 |
| | V10 | 615 | 20 |
| | V11 | 664 | 27 |
| | V12 | 692 | 28 |
| | V13 | 720 | 29 |
| | V14 | 750 | 30 |
| | V15 | 780 | 30 |
| | V16 | 812 | 34 |
| Blue (488nm) | B1 | 508 | 20 |
| | B2 | 525 | 17 |
| | B3 | 542 | 17 |
| | B4 | 581 | 19 |
| | B5 | 598 | 20 |
| | B6 | 615 | 20 |
| | B7 | 661 | 17 |
| | B8 | 679 | 18 |
| | B9 | 697 | 19 |
| | B10 | 717 | 20 |
| | B11 | 738 | 21 |
| | B12 | 760 | 23 |
| | B13 | 783 | 23 |
| | B14 | 812 | 34 |
| Red (635nm) | R1 | 661 | 17 |
| | R2 | 679 | 18 |
| | R3 | 697 | 19 |
| | R4 | 717 | 20 |
| | R5 | 738 | 21 |
| | R6 | 760 | 23 |
| | R7 | 783 | 23 |
| | R8 | 812 | 34 |

**Suppl. Table 2. Cytek Aurora ® configuration.**

| Laser | fluorochrome | peak channel | antibody | | | |
|---|---|---|---|---|---|---|
| | | | human blood leukocytes | Biolegend Compensation Beads | UltraComp eBeads Plus | mouse splenocytes |
| | | | anti-human | | | anti-mouse |
| Violet (405nm) | BV421 | V1 | CD19 | CD19 | CD19 | CD4 |
| | SB436 | V2 | CD31 | CD31 | CD31 | CD31 |
| | eFl450 | V3 | CD3 | CD3 | CD3 | CD11b |
| | BV480 | V5 | CD4 | CD4 | CD4 | CD4 |
| | V500 | V7 | CD16 | CD16 | CD16 | CD45 |
| | SV538 | V7 | CD3 | CD3 | CD3 | B220 |
| | BV570 | V8 | CD11b | CD11b | CD11b | CD11b |
| | BV605 | V10 | CD38 | CD38 | CD38 | CX3CR1 |
| | BV650 | V11 | CD15 | CD15 | CD15 | CX3CR1 |
| | BV711 | V13 | CD3 | CD3 | CD3 | CD8 |
| | BV750 | V14 | CD56 | CD56 | CD56 | TCRb |
| | BV785 | V15 | TLR2 | TLR2 | TLR2 | CD8 |
| Blue (488nm) | BB515 | B1 | CD8 | CD8 | CD8 | CD8 |
| | FITC | B2 | CD11b | CD11b | CD11b | CD4 |
| | Alexa532 | B3 | CD45 | CD45 | CD45 | CD45 |
| | PE | B4 | CX3CR1 | CX3CR1 | CX3CR1 | CD4 |
| | PE/Dazzle594 | B6 | CCR2 | CCR2 | CCR2 | CD11b |
| | PE/Fire640 | B7 | CD45RO | CD45RO | CD45RO | CD11c |
| | PE-Cy5 | B8 | CD19 | CD19 | CD19 | CD19 |
| | BB700 | B9 | CD127 | CD127 | CD127 | CD11c |
| | PerCP-Cy5.5 | B9 | CD14 | CD14 | CD14 | CD4 |
| | PerCP-eFl710 | B10 | CD45RA | CD45RA | CD45RA | CD369 |
| | PE-Cy7 | B13 | CD161 | CD161 | CD161 | CD8 |
| | PE/Fire810 | B14 | CXCR3 | CXCR3 | CXCR3 | CD3 |
| Red (635nm) | APC | R1 | CD14 | CD14 | CD14 | CD4 |
| | Al647 | R2 | CD66b | CD66b | CD66b | CD8 |
| | SparkNIR 685 | R3 | CD45 | CD45 | CD45 | CD8 |
| | Al700 | R4 | CD19 | CD19 | CD19 | CD8 |
| | APC-Cy7 | R7 | CD33 | CD33 | CD33 | CD45 |
| | APC/Fire810 | R8 | CD25 | CD25 | CD25 | Gr1 |

**Suppl. Table 3. Antibodies used in Figure 1 and Supplementary Figure 1**

| Manufacturer | Product name | Catalog # | species reactivity | | | | | |
|---|---|---|---|---|---|---|---|---|
| | | | mouse | rat | rabbit | hamster | human | donkey |
| Biolegend | Compensation Beads | 424601 | x | x | x | x | x | x |
| ThermoFisher Scientific | Ultracomp eBeads Plus | 01-3333-41 | x | x | x | x | x | |
| | Ultracomp eBeads | 01-2222-42 | x | x | | x | | |
| | AbC Total Antibody Compensation Bead kit | A10513 | x | x | x | x | | |
| Beckman Coulter | VersaComp Antibody Capture Kit | B22804 | x | x | x | x | | |
| Cytek Biosciences | Cytek® FSP™ CompBeads | B7-10011 | x | x | | x | | |
| Novus Biologicals | Anti-Mouse Ig (H+L) Comp-Bead 2 Population (3.0-3.4 um) Kit | NBP3-11302 | x | x | | x | | |
| | Anti-Mouse Ig (H+L) Comp-Bead 3 Population (5.5 µm) Kit | NBP3-00497 | x | x | | x | | |
| | Anti-Mouse Ig (H+L) Comp-Bead 3 Population (7.5 µm) Kit | NBP3-00499 | x | x | | x | | |
| | Blank Comp-Bead Particles (negative control) | NBP3-00500 | | | | | | |
| Miltenyi Biotec | MACS®Comp Bead Kit, anti-mouse Igκ | 130-097-900 | x | | | | | |
| | MACS®Comp Bead Kit, anti-human Igκ | 130-104-187 | | | | | x | |
| | MACS®Comp Bead Kit, anti-rat Igκ | 130-107-755 | x | | | | | |
| | MACS®Comp Bead Kit, anti-REA | 130-104-693 | | | | | x | |
| BD Biosciences | BD™ CompBeads Anti-Rat Ig, κ/Negative Control Compensation Particles Set | 552844 | | x | | | | |
| | BD™ CompBeads Anti-Rat and Anti-Hamster Ig κ /Negative Control Compensation Particles Set | 552845 | | x | | x | | |
| | BD™ CompBeads Anti-Mouse Ig, κ/Negative Control Compensation Particles Set | 552843 | x | | | | | |
| | BD™ CompBead Plus Anti-Rat Ig, κ/Negative Control (BSA) Compensation Plus (7.5 µm) Particles Set | 560499 | | x | | | | |
| | BD™ CompBead Plus Anti-Mouse Ig, κ/Negative Control (BSA) Compensation Plus (7.5 µm) Particles Set | 560497 | x | | | | | |

### Suppl. Table 4. commercially available compensation beads

| fluorochrome | Biolegend® Compensation Beads | UltraComp eBeads™ Plus |
|---|---|---|
| BV421 | ✓ | ✓ |
| Super Bright™ 436 | ✓ | ✓ |
| Pacific Blue™ | ✓ | ✓ |
| BV480 | ✓ | X |
| V500 | ✓ | ✓ |
| Spark Violet™ 538 | ✓ | ✓ |
| BV570 | ✓ | ✓ |
| BV605 | ✓ | ✓ |
| BV650 | ✓ | ✓ |
| BV711 | ✓ | ✓ |
| BV750 | ✓ | ✓ |
| BV785 | ✓ | X |
| BB515 | ✓ | ✓ |
| FITC | ✓ | ✓ |
| Alexa Fluor™ 532 | ✓ | ✓ |
| PE | ✓ | ✓ |
| PE/Dazzle™ 594 | ✓ | ✓ |
| PE/Fire™ 640 | ✓ | ✓ |
| PE-Cy5 | ✓ | ✓ |
| BB700 | ✓ | ✓ |
| PerCP-Cy5.5 | X | X |
| PerCP-eFluor™ 710 | ✓ | X |
| PE-Cy7 | ✓ | ✓ |
| PE/Fire™ 810 | ✓ | ✓ |
| APC | ✓ | ✓ |
| Alexa Fluor™ 647 | ✓ | ✓ |
| Spark NIR™ 685 | ✓ | ✓ |
| Alexa Fluor™ 700 | X | X |
| APC-Cy7 | ✓ | ✓ |
| APC/Fire™ 810 | ✓ | ✓ |

**Suppl. Table 5. optimal reference control types for full spectrum cytometry**

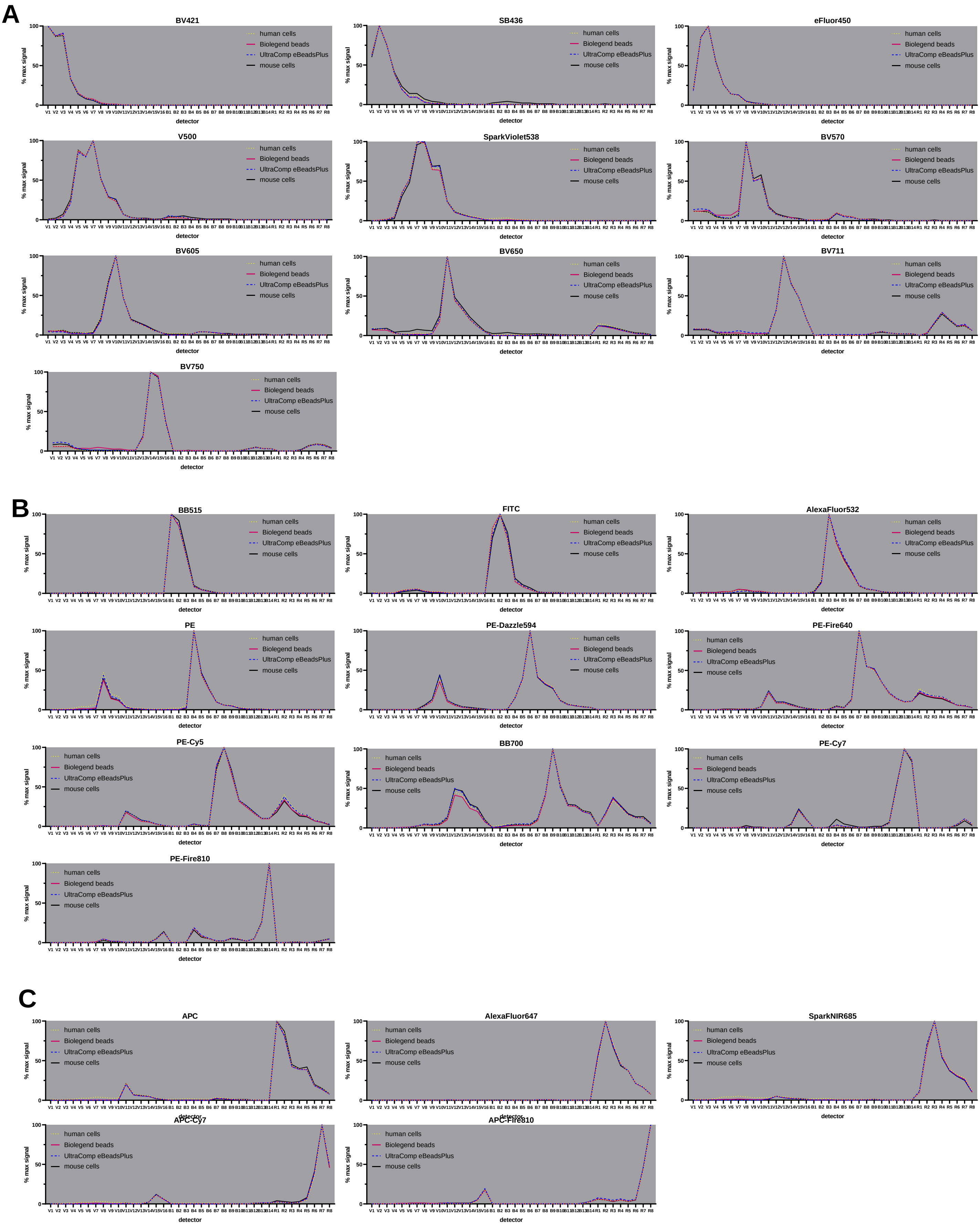


**Supplementary Figure 1. Normalized emission spectra of fluorochromes showing identical emission spectra on cells and beads.** Indicated fluorochromes were stained on cells (human peripheral blood leukocytes or murine splenocytes), Biolegend® Compensation Beads or UltraComp eBeads™ Plus Compensation Beads (ThermoFisher Scientific). Emission spectra were extracted from Spectroflow software, normalized to the maximum signal and overlaid in GraphPad Prism. **A** Fluorochormes excited by the violet laser (405nm). **B** Fluorochormes excited by the blue laser (488nm). **C** Fluorochormes excited by the red laser (635nm).

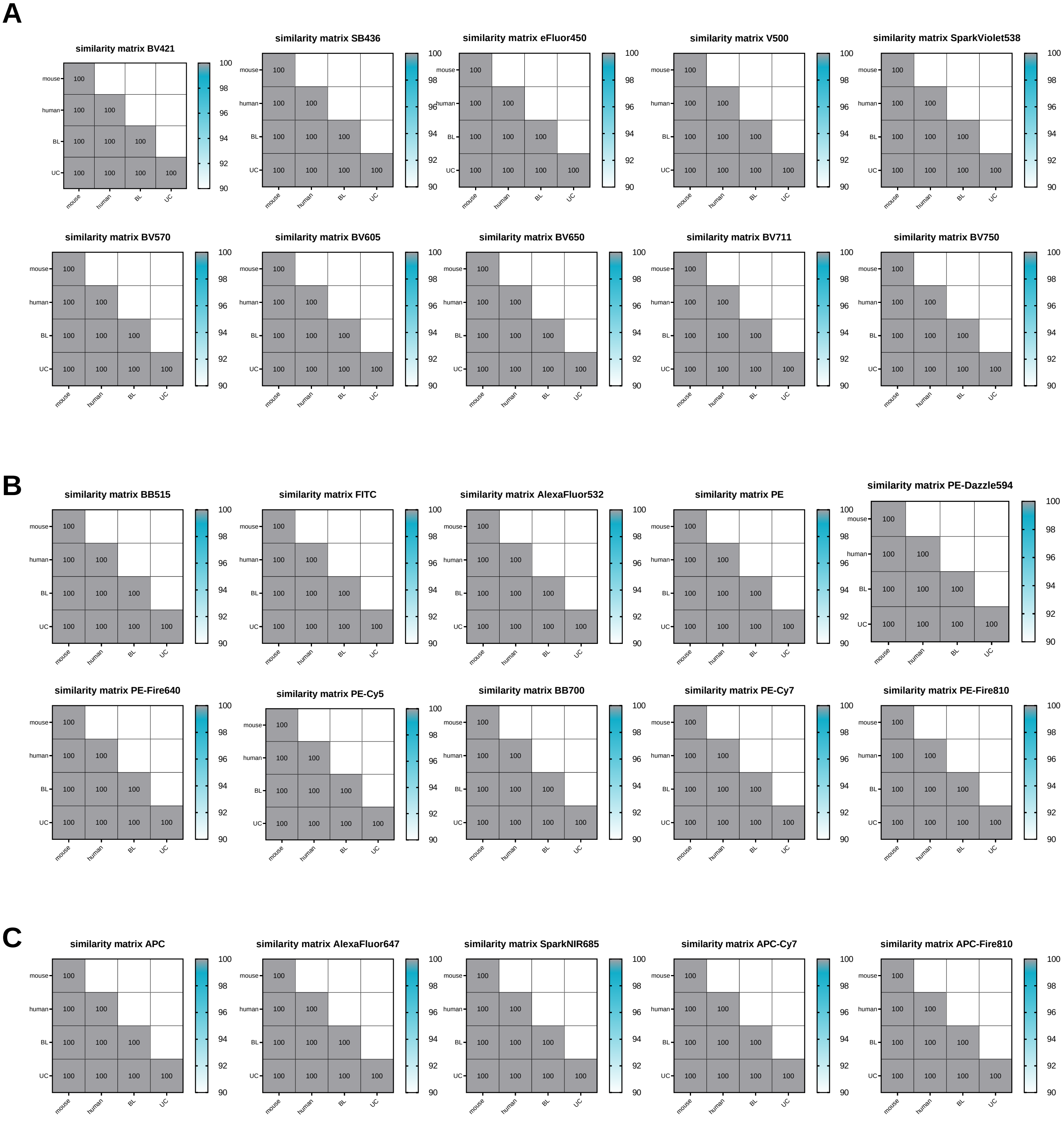


**Supplementary Figure 2. similarity matirzes of fluorochromes showing identical emission spectra on cells and beads.** Indicated fluorochromes were stained on cells (human peripheral blood leukocytes or murine splenocytes), Biolegend® Compensation Beads or UltraComp eBeads™ Plus Compensation Beads (ThermoFisher Scientific). Similarity was determined in Spectroflow software. **A** Fluorochormes excited by the violet laser (405nm). **B** Fluorochormes excited by the blue laser (488nm). **C** Fluorochormes excited by the red laser (635nm). BL= Biolegend compensation beads, human= human peripheral blood leukocytes, mouse= murine splenocytes, UC= UltraComp eBeads Plus

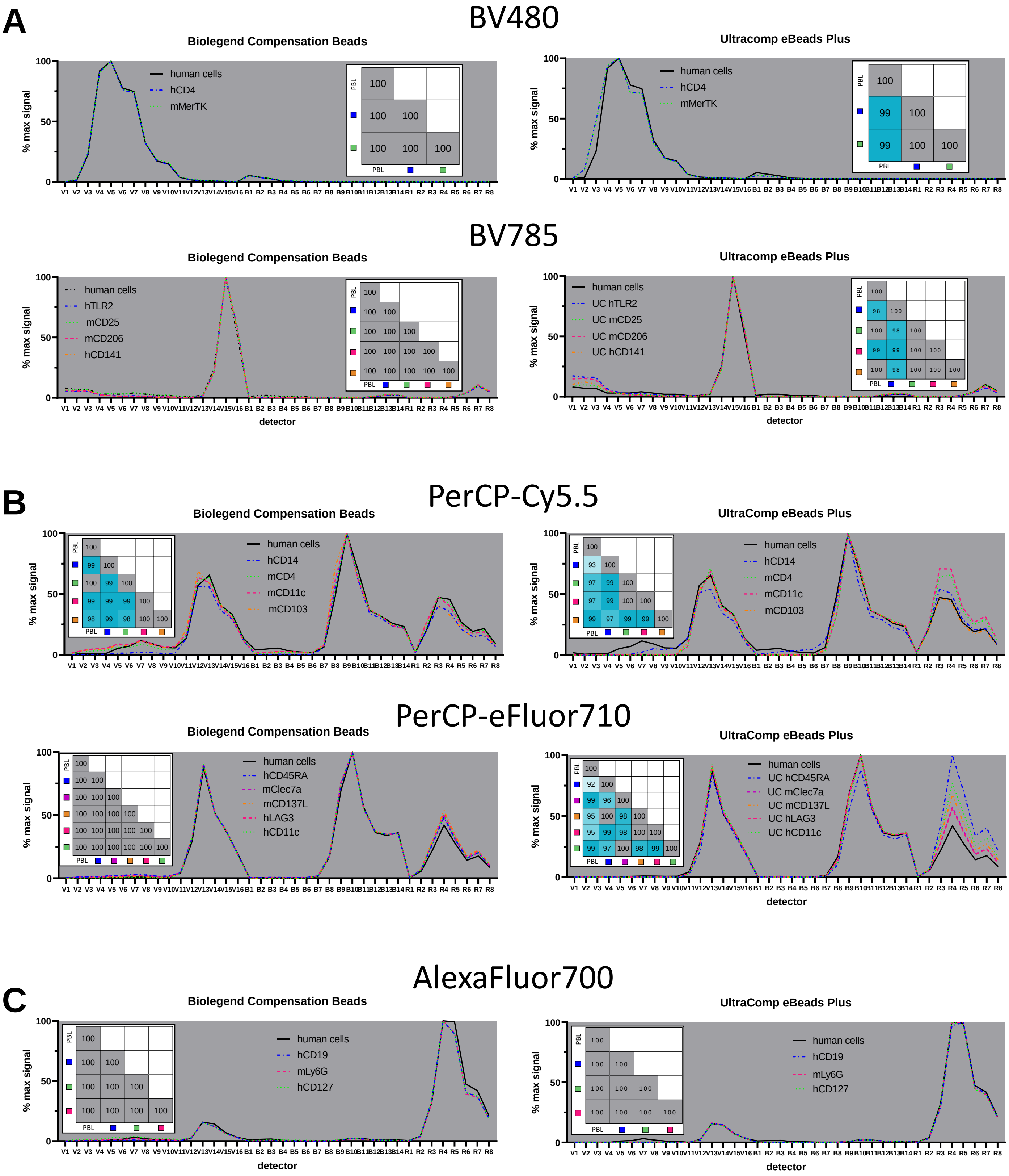


**Supplementary Figure 3.** Indicated fluorochromes were stained on cells (human peripheral blood leukocytes), Biolegend® Compensation Beads or UltraComp eBeads™ Plus Compensation Beads (ThermoFisher Scientific). Emission spectra were extracted from Spectroflow software, normalized to the maximum signal and overlaid in GraphPad Prism. **A** Fluorochormes excited by the violet laser (405nm). **B** Fluorochormes excited by the blue laser (488nm). **C** Fluorochormes excited by the red laser (635nm). h=anti-human, m=anti-mouse, PBL= peripheral blood leukocytes
